# Unsupervised Machine Learning for Species Delimitation, Integrative Taxonomy, and Biodiversity Conservation

**DOI:** 10.1101/2023.06.12.544639

**Authors:** R. Alexander Pyron

## Abstract

Integrative taxonomy combining data from multiple axes of biologically relevant variation is a major recent goal of systematics. Ideally, such taxonomies would be backed by similarly integrative species-delimitation analyses. Yet, most current methods rely solely or primarily on molecular data, with other layers often incorporated only in a *post hoc* qualitative or comparative manner. A major limitation is the difficulty of deriving and implementing quantitative parametric models linking different datasets in a unified ecological and evolutionary framework. Machine Learning methods offer flexibility in this arena by learning high-dimensional associations between observations (e.g., individual specimens) across a wide array of input features (e.g., genetics, geography, environment, and phenotype) to delineate statistical clusters. Here, I implement an unsupervised method using Self-Organizing (or “Kohonen”) Maps (SOMs). Recent extensions called SuperSOMs can integrate an arbitrary number of layers, each of which exerts independent influence on the two-dimensional output clustering via empirically estimated weights. These output clusters can then be delimited into *K* significant units that are interpreted as species or other entities. I show an empirical example in *Desmognathus* salamanders with layers representing alleles, space, climate, and traits. Simulations reveal that the SOM/SuperSOM approach can detect *K=*1, does not over-split, reflects contributions from all layers with signal, and does not allow layer size (e.g., large genetic matrices) to overwhelm other datasets, desirable properties addressing major concerns from previous methods. Finally, I suggest that these and similar methods could integrate conservation-relevant layers such as population trends and human encroachment to delimit management units from an explicitly quantitative framework grounded in the ecology and evolution of species limits and boundaries.

## 1. Introduction

Machine Learning (ML) has had a decades-long impact on ecology and evolution (Azodi et al., 2020; Borowiec et al., 2022; Fountain-Jones et al., 2021; Olden et al., 2008; Recknagel, 2001). These approaches represent part of a fundamental divide in statistics between primarily parametric model-based inference versus algorithmic estimates which treat the data-generating mechanism as unknown (Breiman, 2001). Classical statistics in biology has historically relied on data models, which are typically parameterized based on conceptual understanding of the relevant evolutionary mechanisms and genetic processes. In contrast, the rise of ML methods corresponds with the ascendance of “big” datasets and the recognition that empirical datasets often represent complex, non-linear interactions between variables. The power and flexibility of various ML approaches can easily characterize higher-order patterns in data without constraint from an *a priori* data model that may not include relevant parameters because they are unknown to investigators or difficult to visualize and implement. While this does not obviate the need for pertinent biological mechanisms to draw robust scientific conclusions, it offers the potential for rapid assessment of significant patterns. Such results can then be used for more precise parametric hypothesis-testing and may themselves also reveal complex empirical patterns and potential mechanisms that are nearly invisible to simpler classical approaches.

One area where ML approaches have become particularly relevant is species delimitation (Derkarabetian et al., 2019; Smith and Carstens, 2020). Species delimitation is a complex process involving multiple potential data types, ecological and evolutionary factors, and confounding phylogenetic processes (Burbrink and Ruane, 2021; Carstens et al., 2013).

Concomitantly, integrative taxonomy incorporating multiple lines of evidence is a well-established practice (Dayrat, 2005; Padial et al., 2010), but methods for incorporating molecular (Leaché and Fujita, 2010; Zhang et al., 2013) and morphological (Ezard et al., 2010; Seifert et al., 2014) have rarely been integrated (see Solís-Lemus et al., 2015). A major reason for this is that parametric models linking processes across data types are exceptionally difficult to derive. Consequently, most “integrative” approaches have treated different variables in a more or less *ad hoc* qualitative and comparative fashion instead of quantitative models (e.g., Cicero et al., 2021; Edwards and Knowles, 2014; Fujita et al., 2012; Newton et al., 2020; Pavón-Vázquez et al., 2018; Pyron et al., 2023; Rissler and Apodaca, 2007; Yeates et al., 2011).

Recent ML approaches to species delimitation are also generally not “integrative” in the sense of explicitly combining multiple data sources to produce singular delimitation models. Rather, they can be divided into two categories, both relying primarily on molecular data. First are supervised approaches (Derkarabetian et al., 2022; Perez et al., 2020) such as ‘CLADES’ (Pei et al., 2018) and ‘delimitR’ (Smith and Carstens, 2020). These packages simulate labeled training data under parametric biologically realistic models and train ML classifiers to predict the matching class of generative model for an empirical dataset. One might simulate one-, two-, and three-species models for *n* individuals and *m* molecular loci. Then, an empirical *n*\**m* dataset is fed into the classifier, which predicts that it matches a two-species model with 97% accuracy. Second are unsupervised approaches, which typically utilize some form of dimensionality reduction (e.g., PCA, *t-*SNE, VAE, RF) to extract major axes of genetic variation, then employ some form of clustering such as *k-*means to identify significant divisions (Derkarabetian et al., 2019; Jombart et al., 2010; Pyron et al., 2023). An *n*\**m* dataset might cluster into three distinct segments (*‘K’=*3) representing “species.” Neither type of approach has been extended into a fully “integrative” context where the independent contributions from multiple data types such as alleles, space, climate, and phenotype are estimated jointly.

I present such a method here using Self-Organizing (or “Kohonen”) Maps (SOMs; Kohonen, 1998), expanding the UML species-delimitation approach introduced by Pyron et al. (2023). The original implementation used a single-layer SOM based on a matrix of input features corresponding to allele frequencies at a given set of SNP loci. However, the implementation in the R package ‘kohonen’ (Wehrens and Kruisselbrink, 2018) allows for a “SuperSOM” in which the matrix of input features is divided up into multiple distinct layers, each of which contributes to the output grid (Wehrens and Buydens, 2007). A distance weight is calculated for each layer separately, rescaling contributions from the layers such that differing scales between layers do not overwhelm contributions to the outputs from any one layer, and the relative importance of each layer is estimated directly. I employ this strategy to implement UML species delimitation incorporating data from SNPs, spatial autocorrelation, climate variables, and potentially other features such as phenotypic traits. This approach is implemented in *R* and a tutorial and example files are given at https://github.com/rpyron/delim-SOM.

These SOM-based methods have several unique advantages over all existing approaches. One that we described previously (Pyron et al., 2023) is replicated learning instances to estimate cluster-assignment variability, which produces values analogous to individual ancestry coefficients (e.g., Pritchard et al. 2000; Frichot et al. 2014; Beugin et al. 2018) for each specimen. Both approaches allow for *K=*1 (Cullingham et al., 2020; Janes et al., 2017) and integrate across similarly-explanatory values of *K* (Lawson et al., 2018; Meirmans, 2015). The SuperSOM approach described here adds the capacity to independently integrate multiple data types and estimate their contribution to a unified species-delimitation model (Edwards and Knowles, 2014; Padial and De la Riva, 2021; Schlick-Steiner et al., 2010). While other ML approaches are likely possible and potentially more powerful to be explored in the future, the SuperSOM algorithm offers substantial promise to unify species delimitation and integrative taxonomy. Such a synthesis has been described as desirable by numerous previous authors, but not yet implemented computationally in a tractable framework.

I also echo multiple recent authors in suggesting that ML methods have the potential to radically accelerate progress in species delimitation and integrative taxonomy (Derkarabetian et al., 2019; Smith and Carstens, 2020). A major issue for both parametric and algorithmic approaches has been the presence of multiple competing signals and alternative delimitation models (Derkarabetian et al., 2022; Martin et al., 2021). Yet it has long been recognized that increasing the number and amount of data tends to yield more congruent estimates (Edwards and Knowles, 2014; Newton et al., 2020). Since species are real, discoverable entities in nature (Burbrink et al., 2022; Ghiselin, 1974) explicitly quantitative estimates of their existence should converge as integrative datasets grow in size and dimensionality (de Queiroz, 2007). I term this the “taxonomic convergence hypothesis” and suggest that UML methods such as SuperSOMs can facilitate this systematic unification of species delimitation and integrative taxonomy. This can overcome the potential issue of over-splitting (Parker et al., 2022; Piñeros et al., 2022), and should robust and stable estimates of species boundaries in all but the most difficult cases (O’Connell et al., 2022; Eric Pante et al., 2015).

These methods can also be extended to incorporate conservation-relevant data such as population trends and human encroachment to facilitate delimitation of management units from an explicitly quantitative framework grounded in the ecology and evolution of species limits and boundaries (Crandall et al., 2000; Moritz, 1994; Stanton et al., 2019). Such an approach is easily facilitated by the flexible algorithms implemented here, which allow additional layers containing nearly any type of data. The delimited units may or may not be considered species depending on one’s philosophical viewpoint (Burbrink et al., 2022; Frankham et al., 2012; Karl and Bowen, 1999), but they will nonetheless represent conservation-relevant population segments. Ultimately, the use of ML methods in species delimitation can facilitate not only integrative taxonomy, but better overall integration of ecology, evolution, and conservation.

## 2. Materials & Methods

### 2.1 SuperSOMs

Generally speaking, SOMs are artificial neural networks trained using competitive learning (rather than error correction via back-propagation with gradient descent), assessed by relative distance to the closest unit across clusters (Oja and Kaski, 1999). They can be used produce two-dimensional representations of complex datasets which preserved the higher-dimensional topological structure of the input data. For a dataset with *n* observations of *p* variables, a two-dimensional map represents clusters of individuals assigned to grid cells with similar values across the input features, where nearby clusters have lower multidimensional distances than more distant clusters. These output clusters are projected onto a grid containing a fixed number of cells. This grid can take a variety of shapes and sizes, but I use a simple rectangular grid of 5*sqrt(*n*), a common rule of thumb for appropriate dimensionality reduction using SOMs (Tian et al., 2014). The number of clusters (equivalent to *‘K’* from a STRUCTURE-type analysis) is chosen by *k*-means clustering occupied cells into proximate units based on the weighted sum of squares of neighbor distances between adjacent cells.

The original implementation used a single-layer SOM based on a matrix of input features corresponding to allele frequencies at a given set of SNP loci. However, the implementation in the R package ‘kohonen’ (Wehrens and Kruisselbrink, 2018) allows for a “SuperSOM” in which the matrix of input features is divided up across multiple distinct layers, each of which contributes to the singular output grid (Wehrens and Buydens, 2007). A distance weight is calculated for each layer separately, rescaling contributions from the layers such that differing scales between layers do not overwhelm contributions to the outputs from any one layer, and the relative importance of each layer is estimated directly. I employ this strategy to implement UML species delimitation incorporating data from SNPs, spatial autocorrelation, climate variables, and potentially other features such as traits or phenotypes.

Many default implementations of SOMs have paid little attention to the hyperparameters of neighborhood type (Gaussian, bubble, or heuristic), learning rate (α), and run length (number of steps), finding them to have relatively small impacts on learning outcomes (Pyron et al., 2023; Wehrens and Kruisselbrink, 2018). Given the importance of accurately quantifying layer contributions, I expand on this concern here. Previous studies have found that Gaussian neighborhoods (rather than bubble or heuristic) and linear learning-rates typically yield optimal results under a wide range of conditions (Natita et al., 2016; Stefanovič and Kurasova, 2011). Consequently, I employ these as the default conditions. The layer distance weights (which I term *‘w’*) are calculated automatically to normalize contributions to the final output grid. In practice they vary by orders of magnitude, with important layer like alleles near zero, and large values for unimportant layers. To scale them for comparison, I therefore used sqrt(1/*w*).

The learning rate as implemented in ‘kohonen’ takes an initial and final value for a that declines linearly over ‘rlen’ updates, the run length in steps. Learning strategies other than linear are possible, but generally have little effect (Natita et al., 2016; Stefanovič and Kurasova, 2011)and are not implemented in ‘kohonen.’ Given that the change from α_0_ to α_1_ is relative to the number of steps, the run length typically does not have much of an influence provided that it exceeds whatever initial period is needed to learn the global structure of the data. Therefore, longer runs typically do not yield increased learning accuracy; there is little additional benefit from a longer “burn-in” as in other traditional types of analyses such as MCMC sampling. Nevertheless, I varied rlen and α by orders of magnitude to determine any possible effects, by comparing their output in terms of mean distance to nearest unit across runs for different combinations of each variable. Similar to previous authors (Natita et al., 2016; Stefanovič and Kurasova, 2011), for rlen=100 I tested α_0.1,0.1_, α_0.5,0.1_, α_0.9,0.1_, α_0.9,0.01_, α_0.5,0.01_, and α_0.5,0.5_. For the default of α_0.5,0.1_, I also tested rlen=200 and 1000. These represent a preliminary survey of hyperparameter impact showing relatively little effect for most values (see Results), but should be investigated in depth across a range of empirical datasets.

Finally, I calculated species assignment probabilities from the classification frequency of each individual into the range of sampled *K*, again employing the logic of DAPC (Jombart et al., 2010; see Pyron et al., 2023) to model uncertainty in assigning observed input features to hypothetical delimitation models. In a DNA-only SOM, these are analogous to individual genomic ancestry coefficients (e.g., Beugin et al., 2018; Frichot et al., 2014; Pritchard et al., 2000). In a SuperSOM that reflects the influence of multiple data layers, these are not solely indicative of genetic ancestry and might be better thought of as “species coefficients” which may be useful for a variety of further ecological and evolutionary hypothesis testing. New data can be mapped to an existing SOM, but retraining is fast and will likely represent the simplest solution for updated datasets in most instances rather than attempting to map across replicates.

In a small change from the original implementation (Pyron et al., 2023), I now calculate BIC for the values of *‘K’* in each learning replicate, rather than relying on raw WSS. When *K=*1 (lowest BIC), this value was chosen. Otherwise, the optimal *‘K’* for each run was estimated using the “elbow” method under the ‘diffNgroup’ criterion employed by DAPC (Jombart et al. 2010) from the BIC estimate for grid-cell neighbor distances as described above. I also include 1′BIC using an approximation of the second-order rate of change proposed by Evanno et al. (2005) for use as heuristic to determine when larger values of *K* result in non-zero net improvement, but this is only for comparison and is not used to calculate optimal *K* across runs. Consequently, *K* can therefore also vary across runs, and one can represent this uncertainty in the summarized output. Consequently, each *K* and the estimated ancestries thereof are sampled in rough proportion to their occurrence in the “posterior” distribution of potential learning outcomes for a two-dimensional map. This strategy allows one to incorporate variation in both *K* across runs and the uncertainty in assigning individuals to each of the *K* clusters within runs.

For instance, 100 individuals may cluster strongly into *K*=2 groups (*K_1_* and *K_2_*), each with 50 members. Given potential geographic genetic structure within *K_2_*, an additional 25 members of *K_2_* may sometimes form a third cluster *K_3_* at a low frequency, perhaps 33% of the time. Consequently, we would estimate 100% *K_1_* ancestry for the 50 members of *K_1_*, 100% *K_2_* ancestry for the 25 ‘pure’ members of *K_2_*, and 67%/33% *K_2_*/*K_3_* assignment for the other 25 structured members of *K_2_*. Hybrid ancestry would also be estimated simultaneously, such as other individuals having *F_1_* genotypes from *K_1_*\**K_2_* occurring at ∼50% frequency in each cluster. While possibly confounding different sources of mixed or uncertain ancestry, this approach offers a potentially robust solution to the long-standing problem of representing multiple different but similarly probable estimates of *K* (Cullingham et al., 2020; Lawson et al., 2018). For the problem of label matching across runs, I performed an initial cluster assignment for a range of *K* using DAPC to obtain a fixed reference point, though this was not used in the SOM modeling. After obtaining the SOM replicates, I synchronized labels using an approximate implementation of a non-shortcut CLUMPP-like algorithm (Jakobsson and Rosenberg, 2007) which minimized the distance between the DAPC labels and each SOM replicate.

### 2.2 Re-analysis of Desmognathus

The genetic, spatial, and environmental data come from 71 individuals from 71 sites, while the phenotypic data are expanded to include up to 163 specimens from those sites (Pyron et al., 2023), with the mean of each measurement taken after size correction. The allele frequencies come from a GBS matrix of 5,174 SNPs and 10,526 alleles after trimming to 80% completeness. The space variables are latitude, longitude, and elevation. The climate variables are Level IV Ecoregion (Omernik and Griffith, 2014), HUC4 watershed (Berelson et al., 2004), ENVIREM - monthCountByTemp10 (Title and Bemmels, 2018), and WorldClim - BIO15 (Fick and Hijmans, 2017). The phenotype variables are 17 linear morphometric measurements to 0.01mm precision: SVL (snout-vent length), TL (tail length), AG (axilla-groin length), CW (chest width), FL (femur length [rather than hindlimb length]), HL (humerus length [rather than forelimb length]), SG (snout-gular length), TW(tail width at rear of vent), TO (length of third toe), FI (length of third finger), HW (head width), ED (eye diameter), IN (internarial distance), ES (eye-snout distance), ON (orbito-narial distance), IO (inter-orbital distance), and IC (inter-canthal distance). Here, I size-correct these by SVL using pooled groups (“population2”) in ‘GroupStruct’ (Chan and Grismer, 2022; Onn and Grismer, 2021), then take the mean by site. I also included the larval spot count data for 66 of the 163 individuals from 40 of the sites, taking the mean of left and right counts by site.

For the SuperSOMs including lat/long/elevation (space), environment (climate), and phenotype (traits), I rescaled the input data using minmax normalization to the interval [0,1] for equivalence with the allele frequencies. While this is not strictly necessary given the estimation of layer-specific distance weights, the effects of layer scale are poorly known, and I attempt to avoid this complication by using equivalent units. For each SuperSOM, I performed 100 learning replicates with rlen_100_, α_0.5,0.1_, and a Gaussian neighborhood function. Finally, I compared the individual ancestry coefficients from ‘sNMF’ and the DNA-only SOM (Pyron et al., 2023) with the species coefficients from the SuperSOM. All data and code are given at GitHub repository https://github.com/rpyron/delim-SOM/tree/main.

### 2.3 Simulations

As an initial exploration and demonstration of the capabilities and limitations of this approach, I conducted a few basic simulations to understand commonly asked questions regarding species-delimitation algorithms. First is the possibility or tendency to over-split taxa based on what are perceived to be minor differences. To this end, I simulated a similarly-sized binary SNP matrix (5,174 loci for 71 individuals) for *K=*1 using the R package ‘simulMGF’ (Sikorska et al., 2013). I analyzed this matrix using a DNA-only SOM to ensure the algorithm consistently estimated *K=*1 using minimum BIC as the criterion for cluster selection. I then paired this with the empirical space, climate, and trait data for *Desmognathus monticola.* In the absence of any genetic variation, I assert that most researchers would not consider the significant but qualitatively minor variation in geography, environment, and phenotype to represent two distinct species. Correspondingly, I next estimated a 4-layer trait-based SuperSOM including the null *K=*1 alleles layer in place of the empirical SNP dataset. Finally, I reduced the signal from the null *K=*1 alleles layer to show that the size of the layer itself does not overwhelm contributions from the other, smaller layers. To do so, I divided the null allele frequencies by 1,000 to reduce the overall signal in the layer to ∼0. I then estimated a second trait based SuperSOM including the reduced null allele layer and the three other empirical layers.

## 3. Results

### 3.1 Empirical Delimitation

The estimates from the 4-layer trait based SuperSOM are essentially identical to the previous results from the DNA-only SOM (Pyron et al., 2023), but also reflect significant contributions from space, climate, and traits (Fig. 1, 2). Of the latter three, phenotype has a larger impact than climate, and both are substantially more important than space alone. Converting optimal cluster selection to BIC reveals a broad minimum plateau at *K=*3–7, indicative of potential over-splitting common to such methods (Cullingham et al., 2020; Lawson et al., 2018), but use of a conservative elbow criterion minimizes over-fitting and yields 100% support for *K=*2. Evaluation of learning performance across a broad range of hyperparameter values for learning rate and run length revealed relatively little variation outcomes (Fig. 3). Decreased performance (increased relative distances to closest unit) is observed for very low and very high values (α_0.5,0.01_ and α_0.5,0.5_) of final learning rate, but increasing run length has little effect from 100 to 1,000. Finally, comparison of individual ancestry coefficients from sNMF and the DNA-only SOM reveals a tight linear relationship for hybrids between ∼30–70% ancestry (Fig. 4a). Outside of this range, SOM coefficients are essentially bimodal in predicting ∼100% identity. In contrast, trait based SuperSOM “species coefficients” are essentially binary, predicting dominant group membership at ∼100% identity (Fig. 4b) based on integrated datasets.

**Figure 1.**
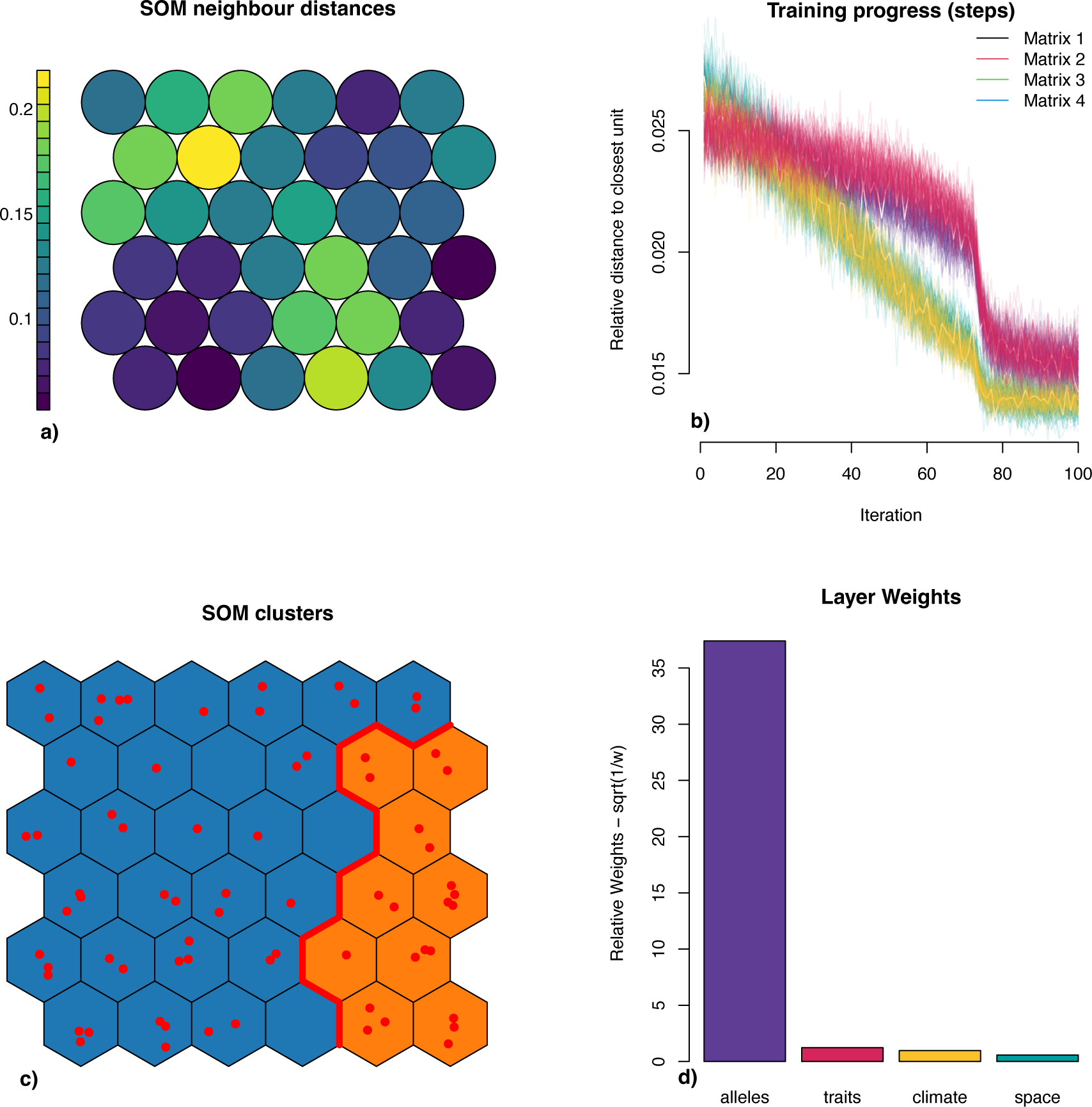
Results from SuperSOM training on the 4-layer *Desmognathus monticola* dataset (alleles, space, climate, traits), showing a) neighbor distances between output grid cells, b) training progress for the four different data layers, c) output grid clustered into the optimal number of clusters, and d) scaled weights for the input data layers.

**Figure 2.**
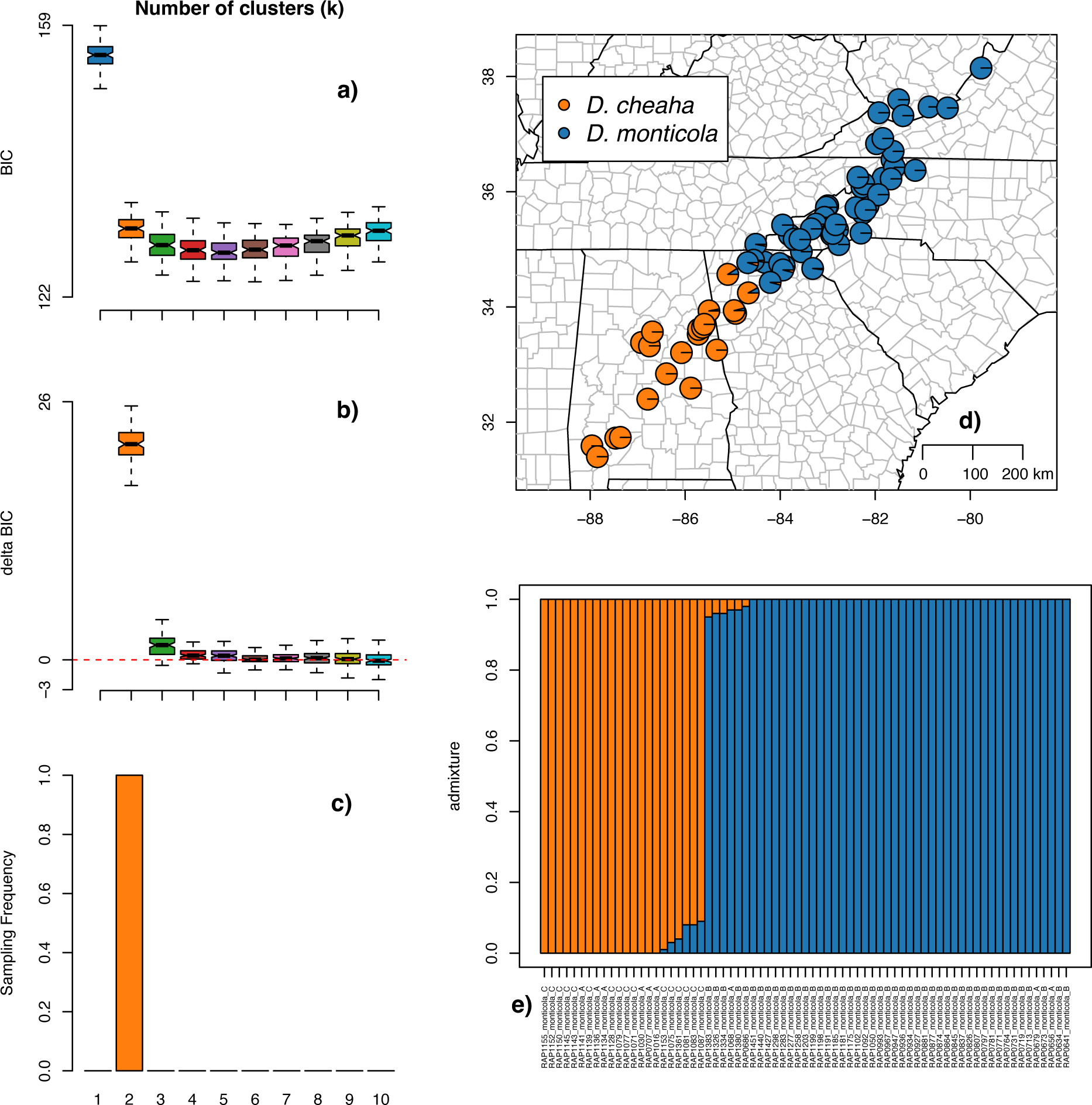
Clustering output from the 4-layer SuperSOM analysis of alleles, space, climate, and traits, showing a) BIC values for 100 replicates across values of *K,* b) second-order ΔBIC, c) sampling frequency for values of *K* based on optimal selection using the elbow criterion, d) a map of specimens with pie charts representing species coefficients of 71 specimens from 71 sites, and e) a STRUCTURE-type barplot of species coefficients.

**Figure 3.**
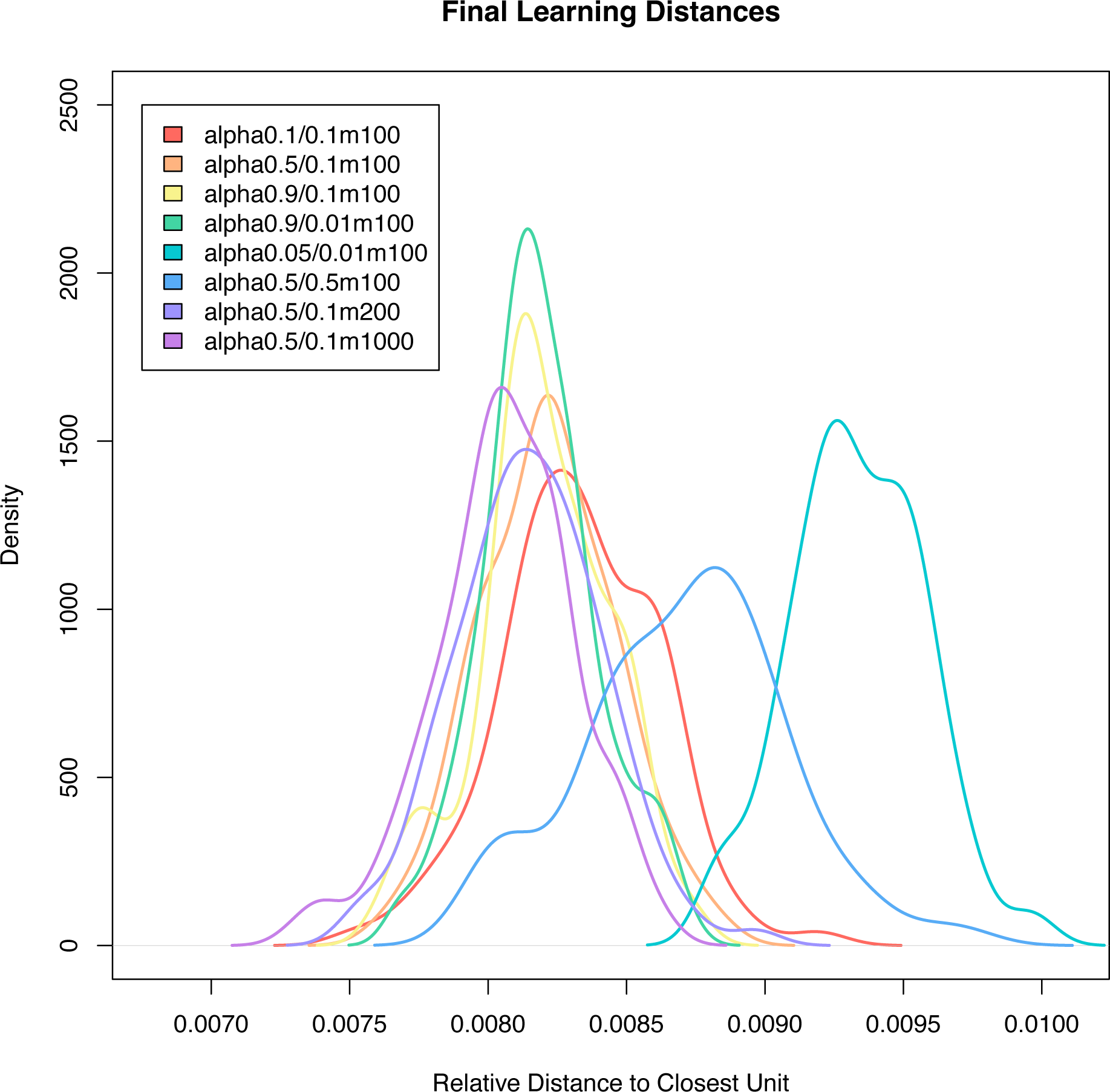
Evaluation of learning performance (relative distance to closest unit) across a range of hyperparameter values for learning rate (α_0_/α_1_) and run length (rlen), showing relatively little impact of rlen and decreased performance only at extreme values of α_1_. The degree to which the scale of these hyperparameters is context-specific is poorly known and should be investigated in more detail (Natita et al., 2016; Stefanovič and Kurasova, 2011).

**Figure 4.**
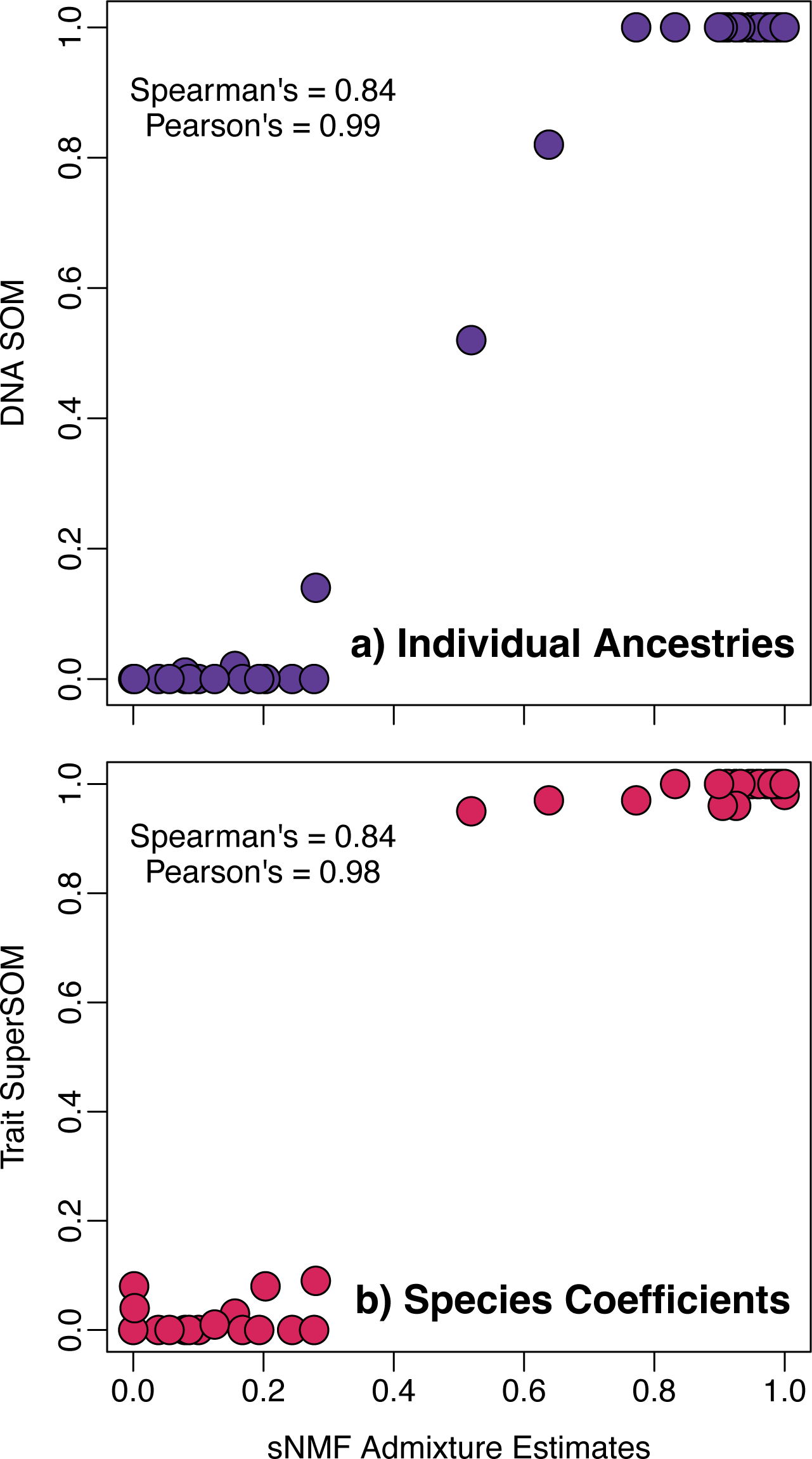
Comparison of a) individual ancestry coefficients from a DNA-only SOM and b) species coefficients from a 4-layer SuperSOM with layers representing alleles, space, climate, and traits to the admixture estimates from sNMF given by (Pyron et al., 2023). Relationships between genetic hybrid indices are roughly linear between ∼30–70% (a), but species coefficients are essentially binary for the two species when considering additional axes of geographic, environmental, and phenotypic variation across genomic ancestry proportions (b).

### 3.2 Simulated performance

The simulations reveal several desirable properties for a species-delimitation method in comparison with existing approaches. First, conversion to optimal cluster selection using BIC yields strong support for *K=*1 when no genetic structure is present, a major advantage over many previous methods (Cullingham et al., 2020; Evanno et al., 2005; Janes et al., 2017). For the simulated *K=*1 DNA-only SOM, *K=*1 was selected in all replicates and the second-order 1′BIC for higher values did not differ from 0 (Fig. 5a–c). Second, this advantage is maintained even when paired with significant geographic, environmental, and phenotypic structure that would not otherwise likely be indicative of speciation (Fig. 5d–f). Third, the size of the alleles layer alone does not necessarily overwhelm the contribution of other layers when relatively little genetic divergence is present. The 4-layer trait based SuperSOM including the null alleles layer estimated *K=*2–5 with *K_2_=*52% of samples and the highest contributions again coming from traits, climate, and space, with alleles having the smallest weight (Fig. 5g–l).

**Figure 5.**
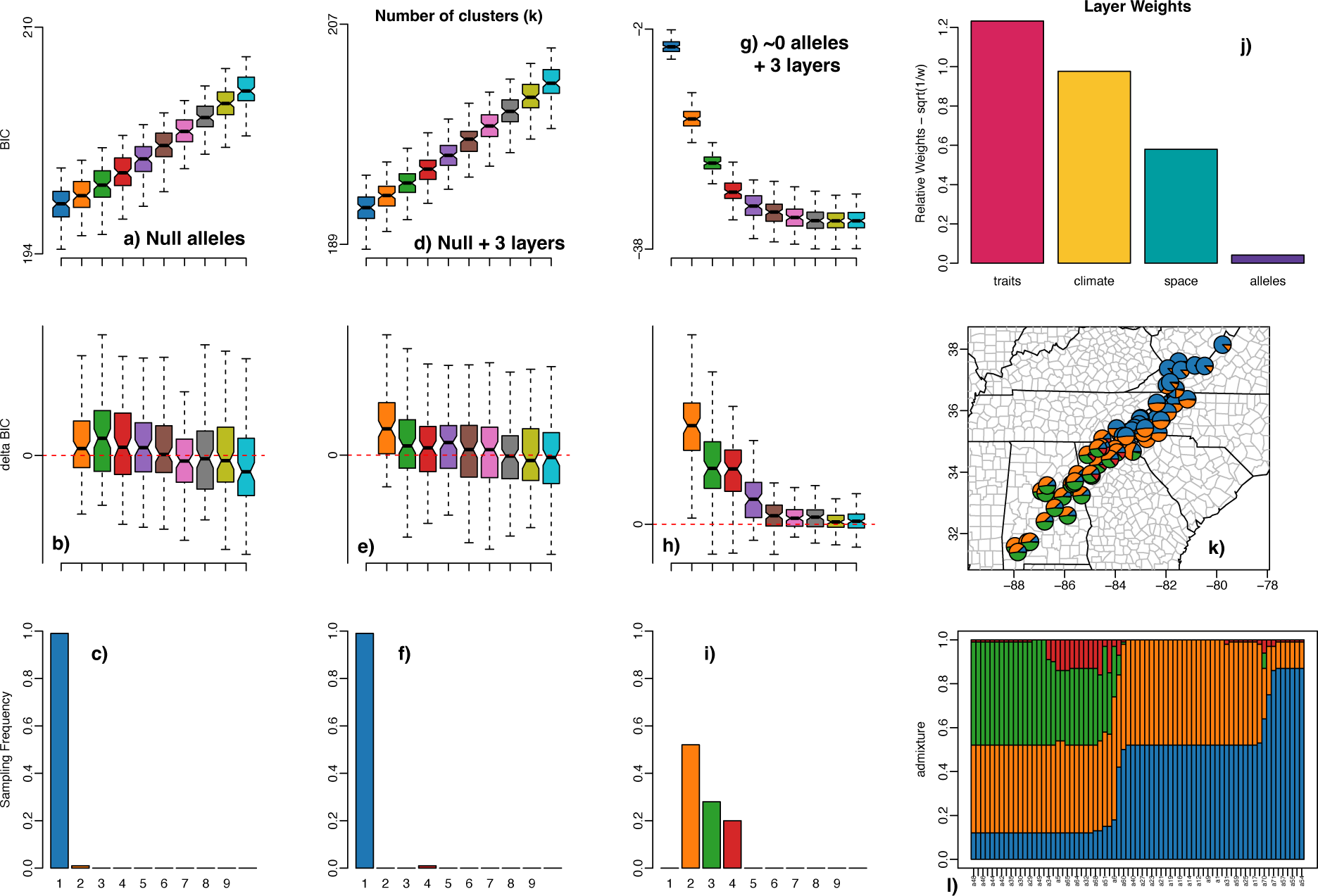
Results of simulation analyses showing overwhelming support for *K=*1 when only a single genetic population is present (null alleles; a–b) and when these null alleles are combined with the empirical space, climate, and trait layers (d–f). Reducing the signal in the null alleles to ∼0 supports *K_2-4_* (g–l), reflecting the signal present in geographic, environmental, and phenotypic layers alone (j), which still estimate a roughly binary division corresponding to *D. cheaha* in the south and *D. monticola* in the north (k), sampling multiple possible ancestries (l).

## 4. Discussion

### 4.1 UML delimitation using SuperSOMs

Simulations reveal that the SOM/SuperSOM approach can detect *K=*1, does not over-split, reflects contributions from all layers with signal, and does not allow layer size (e.g., large-scale genetic matrices) to overwhelm other datasets. These have been major concerns for many previous researchers addressing the problem of species delimitation, particularly for algorithmic methods relying solely or primarily on genetic data (Luo et al., 2018). The method proposed here has several major advantages over existing approaches. As with most ML algorithms, it is extremely flexible and can be implemented over a wide variety of data types and formats without the limits imposed by many formal models of phenotypic and DNA-sequence evolution (Solís-Lemus et al., 2015; Sukumaran et al., 2021). Additionally, I have implemented it to use the four data layers most likely to be commonly available in empirical questions (alleles, space, climate, and traits), which is essentially unique among species-delimitation methods implemented to date.

When estimating species-delimitation models from a DNA-only SOM, the estimated species coefficients are essentially analogous to individual ancestry coefficients, admixture estimates, or hybrid indices from methods such as STRUCTURE, ADMIXTURE, or sNMF (Alexander et al., 2009; Frichot et al., 2014; Pritchard et al., 2000). The obvious difference is that they are not based on an underlying genetic model. Consequently, the empirical values here tend to exhibit a more sigmoidal relationship with the sNMF estimates; outside of ∼30-70% mixed ancestry, SOMs predict ∼100% parental identity given that there is very little uncertainty in cluster assignment in the tails. SuperSOMs heighten this effect, producing nearly binary species coefficients for most populations. This is unsurprising given that even admixed populations near the hybrid zone tend to be either montane or Piedmont and have the strongly diagnostic character of 4–5 versus 6–7 larval spots in *D. monticola* compared to *D. cheaha* (Pyron et al., 2023). I do note that the sample size here is relatively small, as individual ancestry coefficients were already sharply bimodal with relatively few hybrid or admixed individuals.

The approach described here is also robust to a wide range of hyperparameter values, suggesting that learning performance will be trustworthy across a range of empirical conditions. Similar values have been implemented across a variety of smaller datasets and yield similarly congruent results (Natita et al., 2016; Stefanovič and Kurasova, 2011), although the fine-scale impact of grid topologies, learning schedules, and neighborhood types on species delimitation in particular should be addressed in more detail in future research. While further investigation of empirical performance across a range of conditions is warranted, the relatively simple nature of SOMs makes them an ideal candidate for an easy to use, flexible, and generally applicable method while alleviating concerns about performance related to complex hyperparameters.

### 4.2 The “taxonomic convergence hypothesis”

A major issue in systematics is the often still lingering separation between species delimitation and taxonomy; species need to be described, not just discovered and delimited (E. Pante et al., 2015). While the operationalization and implementation of “integrative” taxonomy is still debated (Conix, 2018), the notion itself is increasingly widespread (Vinarski, 2020; Zamani et al., 2022). Although there are many contributing factors to the taxonomic impediment (Engel et al., 2021), one set of three in particular is specific to integrative taxonomy. The first is the concept of taxonomic inflation, particularly when relying on limited molecular datasets and phylogenetic delimitation (Zachos et al., 2013). Yet, numerous recent studies demonstrate the potential for taxonomic reduction from integrated datasets (Freitas et al., 2020; Hundsdoerfer et al., 2019; Parker et al., 2022; Piñeros et al., 2022; Pyron et al., 2016). Second is the concern that common evolutionary models will over-split species based on population-level divergence (Sukumaran and Knowles, 2017). Yet, while datasets will almost always predominantly reflect signal from large multi-locus or genome-scale markers (Fujita et al., 2012), there is no inherent limit on the amount of informative environmental and phenotypic data (or behavior, physiology, ecological interaction, etc.) that can be gathered for most taxa.

Third is the more durable issue of disagreement between various species-delimitation methods which often induce hesitancy or indecision for researchers (Carstens et al., 2013). These maybe minor (e.g., support for 6 versus 7 species), or represent major conflict or contradictory outcomes. In contrast to concerns of “inflation” or “over-splitting” which are often easily addressed by integrative analyses in numerous taxonomic groups, the problem of congruence has not been widely solved. Yet, I predict it will abate in the future for three major reasons. One is that datasets continue to grow; historically, conflicts have often arisen when sampling only a few loci and traits. Two, a great deal of conflict has arisen from methods employing different strict criteria, thresholds, or models for delimiting species based on barcode gaps, branch lengths, or coalescent units (Puillandre et al., 2012; Yang and Rannala, 2014; Zhang et al., 2013). Three, few if any quantitative methods to date have actually incorporated a wide range of biologically relevant factors other than genes and phenotypes such as space and environment, despite frequent calls for such approaches (Rissler and Apodaca, 2007; Solís-Lemus et al., 2015).

Increasing the size and comprehensiveness of datasets and flexibility of methods addresses all three naturally as a consequence of greater empirical and methodological integration.

Consequently, I propose the “taxonomic convergence hypothesis.” For most groups that have a well-characterized natural history, integrative taxonomic approaches that combine multiple data types across a broad range of biologically relevant axes of variation will eventually tend to yield congruent delimitation models as datasets grow in size. The use of ML approaches such as the SuperSOM algorithm implemented here will likely help to facilitate and accelerate that convergence. Situations where the TCH may not hold include poorly known taxa and regions (e.g., under-explored tropical biomes, the deep sea) for which such data are unavailable (Cordier et al., 2022; Goodwin et al., 2015), “dark” taxa and other groups (e.g., microbes) that are difficult to characterize by ecology and phenotype (Page, 2016; Sanford et al., 2021), and complex hybrid interactions that present misleading genotypic evidence (Chan et al., 2021) or strain the concept of “species” itself (Bogart, 2019).

### 4.3 Directions for future research

The third factor listed above (complex genomic ancestry) is perhaps the most immediately relevant for many groups and of concern to a wide variety of researchers. What happens with hybrids? A tendency for admixed genotypes to emerge as distinct clusters is a known problem for a variety of phylogenetic and species-delimitation methods (Chan et al., 2021; Dolinay et al., 2021), and has even been observed in other *Desmognathus* species (Pyron et al., 2022). While known from several empirical cases, the behavior of most methods in the face of such complexities has not been investigated in detail and is not well understood. It seems likely that UML approaches such as SuperSOMs and other algorithms (Derkarabetian et al., 2019) will exhibit similarly poor or aberrant performance in such instances. Parametric genetic approaches may address this issue by implementing sufficiently parameterized models to account for complex genomic ancestry. Similarly, ML algorithms might merge supervised and unsupervised approaches (Derkarabetian et al., 2022; Smith and Carstens, 2020) by pre-training classifiers to recognize the allelic signature of hybrid genotypes arising from various processes. These methods are still in their infancy, and there is an open field to characterize their behavior under a wide range of empirically relevant biological conditions and develop them further.

One potentially positive synergy that remains almost totally unexplored is the potential integration of data regarding threat and risk for conservation management (Crandall et al., 2000; Giangrande, 2003; Mace, 2004; Moritz, 1994). It is well known that accurate and precise species delimitation is extremely relevant to biodiversity conservation (Shirley et al., 2014; Stanton et al., 2019). Simultaneously, numerous recent authors have suggested considering conservation in taxonomic practice (Conix, 2019; Frankham et al., 2012). Yet, few if any studies have directly incorporated conservation-relevant data into species-delimitation analyses. This primarily stems from the lack of any integrative quantitative methods for handling such analyses but approaches such as the SuperSOM algorithm implemented here could easily be extended to incorporate a fifth “conservation” layer containing data such as population trends and human encroachment.

Many researchers (myself included) reject the notion that the concept of “species” as historical evolutionary entities should be altered to incorporate contemporaneous threats and risks (Burbrink et al., 2022; Karl and Bowen, 1999; Pyron and Mooers, 2022; Russello and Amato, 2014). However, this does not preclude using species-delimitation algorithms to delimit population segments for conservation managements based not only on their genetic, geographic, environmental, and phenotypic distinctiveness, but also on relevant indicators such as population trends and measurements of risk and threat such as human encroachment and sensitivity to climate change or habitat alteration. For example, including data on recent changes in abundance and habitat fragmentation would likely reveal the physiographically disjunct “Red Hills” populations of *Desmognathus cheaha* to be a distinct unit. These populations are genetically and phenotypically distinct while definitively falling short of “species” status (Casey, 2002; Pyron et al., 2023), but are ecogeographically separate and have experienced significant anthropogenic degradation and local extirpation compared to Piedmont and Appalachian populations (Holzheuser and Means, 2021). Methods for delimiting “species,” particularly flexible ML methods such as the SuperSOM approach introduced here, could easily be co-opted or extended for quantitative analysis of conservation units in an ecological and evolutionary framework guided by taxonomic principles and grounded within species limits and boundaries.

## 5. Conclusions

The SuperSOM method implemented here has several desirable properties, including the ability to detect *K=*1 when population structure is absent, a tendency not to over-split even when larger values of *K* have partial support, reflection of contributions from all layers with significant signal, and control for layer size (e.g., large-scale genetic matrices) not to overwhelm other datasets. This represents a substantial advance in the potential for integrative species delimitation leading to integrative taxonomy by merging multiple biologically relevant axes of ecological and evolutionary variation into a single unified analysis. While challenges remain for taxa and areas that lack such data, groups that can’t be easily quantified beyond genetics, and complex patterns of genomic ancestry including hybridization, future research can extend these methods and address such concerns. One possibility is merging supervised and unsupervised approaches, such as pre-training classifiers on “good” species and hybrid genotypes to recognize such instances. Furthermore, I suggest that this and similar approaches can easily be extended into the realm of biodiversity conservation by including relevant data layers such as population trends and human encroachment to delimit significant management units. While many systematists (myself included) would not treat these as “species,” the conceptual merger of biodiversity conservation and species delimitation will allow for quantitative analysis of conservation units in a taxonomic framework grounded in empirical estimation of species limits and extinction risk.

## Acknowledgments

This research was supported in part by GW UFF awards and US NSF grants DBI-0905765, DEB-1441719, and DEB-1655737. This work was completed in part with resources provided by the High Performance Computing Cluster at The George Washington University, Information Technology, Research Technology Services. Thanks to G. Bradburd (UMich) and R. Wehrens (WUR) for assistance with theory and algorithms.

## Notes

https://github.com/rpyron/delim-SOM

### Competing Interest Statement

The authors have declared no competing interest.

